# Brf1-Mediated RNA Polymerase III Activity Limits Murine Gammaherpesvirus Spread

**DOI:** 10.64898/2026.09.14.751424

**Authors:** K Rapchak, LS Nyberg, LA Johnson, RA Elbert, CJ Huff, ZA Grissom, SE Dremel, JM Tucker

## Abstract

RNA polymerase III (Pol III) activity is upregulated during herpesvirus infection, yet the functional consequences of this response remain poorly understood. To investigate the role of host Pol III transcription during murine gammaherpesvirus 68 (MHV68) infection, we depleted the Pol III transcription factor Brf1, an essential component of the TFIIIB complex required for transcription from Type I and Type II Pol III promoters. Brf1 depletion enhanced MHV68 replication during low multiplicity of infection (MOI) conditions, resulting in increased viral gene expression, viral protein accumulation, infectious virion production, and extracellular viral genome copies. These effects were confirmed to be Brf1-specific through rescue with an siRNA-resistant Brf1 construct. In contrast, Brf1 depletion produced only modest effects during high-MOI infection, suggesting that Brf1-dependent antiviral activity is most important during multistep viral spread. Transcriptomic analysis revealed accelerated induction of interferon-responsive genes early during infection in Brf1-deficient cells, followed by enhanced host transcript depletion at later stages, consistent with amplified host shutoff. Genetic disruption of the RIG-I/MAVS signaling pathway failed to abolish Brf1 antiviral activity, indicating that this phenotype is independent of MAVS-dependent interferon signaling. Finally, plaque assays demonstrated that Brf1-deficient cells supported larger plaques and increased plaque numbers, suggesting enhanced viral spread and/or entry. Together, our results reveal an unexpected role for Brf1-dependent Pol III activity in controlling gammaherpesvirus spread and suggest that virus-induced Pol III activation contributes to host antiviral defense.

**Importance:** RNA polymerase III (Pol III) activity is stimulated by DNA tumor virus infection, yet the biological significance of this response has remained unclear. Utilizing the murine gammaherpesvirus, MHV68, we demonstrate that the Pol III transcription factor Brf1 functions as a previously unrecognized antiviral restriction factor. Depletion of Brf1 enhanced viral gene expression, protein accumulation, infectious virion production, plaque formation, and viral spread. Genetic rescue was able to restore restriction of infection. Surprisingly, this phenotype was most pronounced during low-multiplicity infections and was associated with accelerated viral cell-cell spread. Transcriptomic analysis revealed that Brf1 influences host antiviral responses and the dynamics of virus-induced host shutoff, highlighting an unexpected connection between Pol III transcription and cellular defense pathways. Importantly, Brf1-mediated restriction occurred independently of the canonical RIG-I/MAVS signaling axis. These findings provide new insight into why Pol III transcription is broadly induced during herpesvirus infection and identify Brf1-dependent Pol III activity as a key host mechanism that limits gammaherpesvirus spread.

## Introduction

Transcriptional regulation often dictates the outcome of host-pathogen interactions. Viruses hijack host machinery to synthesize viral transcripts while concurrently dampening normal host transcriptional activity. Previous studies of transcriptional regulation during viral infection have focused on RNA Polymerase II, which synthesizes protein-coding messenger RNAs (mRNAs) made by the virus and the host^1–7^. In contrast, regulation of RNA polymerase III (Pol III) and resulting non-coding RNA transcripts (ncRNAs) is less understood. Pol III-transcribed ncRNAs have roles in a variety of essential processes such as translation (e.g., transfer RNAs)^8^, ribosomal biogenesis (e.g., 5S)^9^, or mRNA splicing (e.g., U6)^10^. These processes are also important to sustain viral replication, yet it remains unclear how Pol III is regulated, how the levels of host ncRNAs change, and how these alterations functionally impact infection.

Pol III activity has been found to be upregulated during infection of multiple herpesviruses^11^. Changes in Pol III output seem to differ depending on the virus and/or infected cell type, with elevated levels of distinct Pol III transcripts reported. For example, HIV-1 infection enhances U6 snRNA levels, EBV infected cells show upregulation of brain cytoplasmic 200 (BC200), and KSHV infection leads to an increase in a non-coding RNA involved in DUSP11 suppression, nc886^12–15^. Reported mechanisms driving Pol III upregulation include Pol II/III crosstalk, increased expression of Pol III recruitment factors, increased binding of Pol III to tRNA gene promoters, and altered chromatin states around tRNA genes^16–20^. Additionally, Pol III transcript processing or protein binding can be dysregulated during infection, triggering cytoplasmic RNA sensors due to accessible 5’ triphosphate ends and/or double stranded structural features^21, 22^. While the mechanism of action for each viral infection and/or cell type may differ, we hypothesize that upregulation of Pol III and the conservation of this response play an important role during DNA virus infection.

Our previous work sought to understand if and how Pol III is upregulated during lytic replication of murine gammaherpesvirus 68 (MHV68) — a mouse model gammaherpesvirus genetically similar to both Kaposi Sarcoma Associated Herpesvirus (KSHV) and Epstein-Barr Virus (EBV) ^17, 20, 23^. MHV68 infection was shown to induce B2 SINE transcripts (^24–26^), which are Pol III genes ancestrally derived from a murine tRNA-Ser^27^. We subsequently explored canonical tRNA expression during MHV68 infection and observed increased premature-tRNA (pre-tRNA) levels from ∼14% of total tRNA genes^17^. Here, we sought to test whether these observed increases in RNA polymerase III transcripts serve a functional role during MHV68 infection.

Pol III relies on three unique promoter types for recruitment to genes for nascent transcription. RNA polymerase III promoters are classified into Type I (5S rRNA genes), Type II (e.g., tRNA and B2 SINEs), and Type III promoters (e.g., U6 snRNA and 7SK genes) based on the unique assembly of transcription factors required for each promoter type. A key transcription factor required for recruitment of Pol III, TFIIIB, is composed of Tbp, Bdp1, and either Brf1 (in the case of Type I and II promoters) or Brf2 (Type III promoters). As both B2 SINEs and tRNAs use Type II promoters requiring Brf1 for transcription, we used Brf1 knockdown as a tool to limit these transcripts during infection. Interestingly, we found that loss of Brf1-dependent RNA polymerase activity led to increases in viral gene expression, protein accumulation, and titers across a multistep infection. Surprisingly, enhanced replication was associated with a more rapid interferon response at the onset of the low MOI infection, but this was counteracted by robust host shutoff as the virus spread more efficiently through the Brf1-depleted culture. Moreover, plaque size assays indicate that viral spread is enhanced in Brf1-deficient cells. Altogether, we provide evidence that Brf1-dependent RNA polymerase III activity plays an anti-viral role against MHV68 infection by limiting viral spread.

## MATERIALS AND METHODS

### Cell line creation and culturing

NIH 3T3 murine fibroblasts (ATCC CRL-1658) and NIH 3T12 murine fibroblasts (ATCC CCL-164) were cultured in high-glucose Dulbecco’s modified Eagle’s medium (DMEM; Gibco 11965092) supplemented with 10% fetal bovine serum (FBS; Gibco 26140079). Cells were maintained antibiotic-free and routinely screened for mycoplasma (Invivogen). NIH 3T3 cells (NIH/Swiss-derived) were used for all genetic manipulation experiments and NIH 3T12s were used exclusively for virus propagation. Knockout cell lines were made by transduction of NIH 3T3 carrying Cas9 with lentiviral vectors derived from pMCB320 (Addgene plasmid #89359; a gift from Michael Bassick).

Overexpression cell lines were made by transduction with lentiviral vectors derived from pLJM1 (Addgene plasmid 91980; a gift from Joshua Mendell). Oligonucleotide sequences for knockdown are in Table 1. pLJM1 derivatives described in this work have been deposited to Addgene. Lentiviruses were packaged in HEK293T cells using third-generation packaging plasmids and were used to transduce NIH 3T3 cells. Knockout cell lines were selected with 2 µg/mL puromycin and 5 µg/mL blasticidin.

**Table 1.**
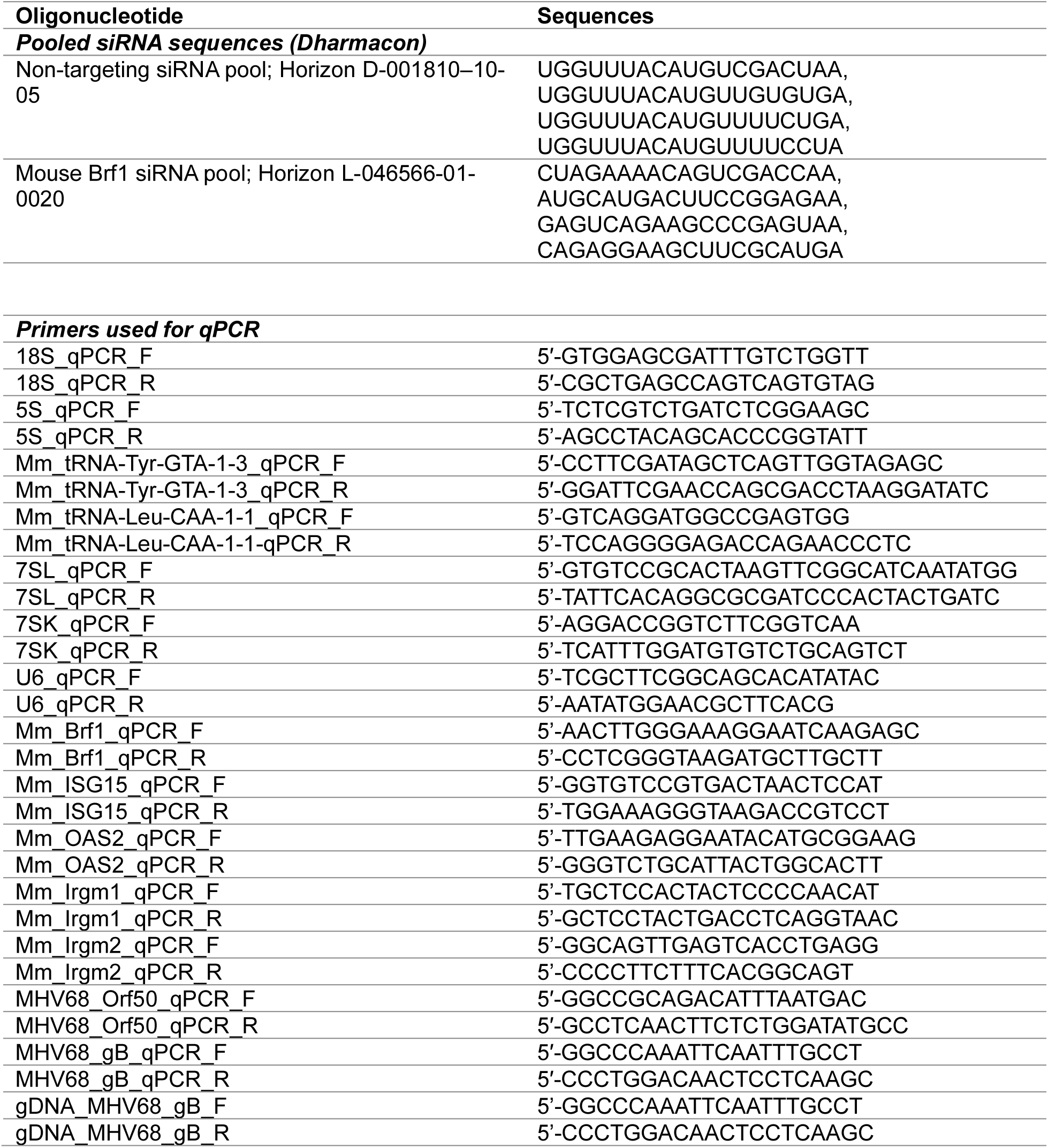
Oligonucleotide sequences.

Overexpression cell lines were selected with 800 µg/mL zeocin.

### Short-Interfering RNA transfection and nucleofection

Pooled siRNAs (sequences in Table 1) were purchased from Dharmacon. For knockdown and overexpression experiments, 1.5 x 10^5^ cells of each respective cell line were seeded into a 6-well culture plate 24 h prior to transfection. Cells were then transfected with siRNAs at a final concentration of 60 nM using RNAiMAX (Invitrogen 13778150). For plaque size experiments, cells were nucleofected using the Neon Transfection System (Thermo Fisher MPK5000). A total of 5.0 x 10^5^ NIH 3T3 cells/well were electroporated with 200 nM siRNAs using the following parameters: 1,400 volts, 20 ms, 2 pulses, and subsequently plated into 6-well plates. Knockdown efficiency for transfection versus nucleofection was determined to be equivalent.

### MHV68 infection

Green fluorescent protein (GFP)-expressing MHV68-MR mutant revertant was propagated in NIH 3T12 cells and used as wild-type virus for this study^28^. To prepare viral stocks, MHV68-infected NIH 3T12 cells and associated culture medium (DMEM + 10% FBS) were subjected to two freeze-thaw cycles. The cellular material was pelleted and discarded. Viruses present in the supernatant were pelleted at 12,000 x g for 2 h and subsequently treated with DNase (Fisher EN0523) for 1 h at 37°C. Following DNase treatment, additional DMEM was added, and virus was pelleted again at 12,000 x g for 2 h. The final virus pellet was resuspended in DMEM + 10% FBS, aliquoted, and stored at-80°C. The titer of the MHV68-GFP stock was determined in NIH 3T3 cells using the 50% tissue culture infective dose (TCID50) assay. For infections, MHV68 was diluted into half the culture volume of DMEM without FBS and applied to corresponding cell lines as indicated. Cells were incubated with virus for 1.5 h at a multiplicity of infection (MOI) of 0.05 or 5. Following incubation, inoculum was removed, cells were washed twice with PBS and replaced with fresh DMEM + 10% FBS.

### Protein quantification and western blots

Cells were washed once with Dulbecco’s phosphate-buffered saline (DPBS; Gibco 14190144) and lysed in radio immunoprecipitation assay (RIPA) buffer supplemented with protease inhibitor tablets (Thermo Fisher 89900, Sigma Aldrich 11836170001).

Lysates were spun by centrifugation at 15,000 rpm for 5 min at room temperature to remove insoluble material. Total protein was denatured in a 4X Laemmli sample buffer containing 2-mercaptoethanol (BME) (Bio-Rad 1610747, Bio-Rad 1610710). For western blot analysis, 30 µg of whole-cell lysate was resolved on 4-15% Mini-PROTEAN TGX gels (Bio-Rad 4561084). Proteins were transferred to polyvinylidene fluoride (PVDF) membranes (Bio-Rad 12023954) using the Trans-Blot Turbo Transfer System (Bio-Rad 1704150). Membranes were blocked for 1 h at room temperature in 5% milk in TBST (Milk: Bio-Rad 1706404, 10X TBS: Bio-Rad 1706435, Tween-20: Sigma-Aldrich P2287-500ML). Blots were incubated overnight at 4°C with primary antibodies (listed in Table 2) diluted in 5% milk/TBST. Membranes were washed in TBST at room temperature and then incubated for 1 h at room temperature with the appropriate horseradish peroxidase (HRP)-conjugated secondary antibodies (listed in Table 2) for 1 h at room temperature. After washing, membranes were treated with Clarity Western ECL or Clarity Western ECL Max substrates (Bio-Rad 1705061, Bio-Rad 1705062) for 5 min and imaged using a ChemiDoc Imager (Bio-Rad 12003153). Band intensity area was quantified using FIJI, and values were normalized to the loading control, GAPDH.

**Table 2.** Antibodies.

| **Target** | **Name, Catalog #** | **Manufacturer, Reported Concentration** | **Host, Clonality, Dilution Used** |
| --- | --- | --- | --- |
| TFIIIB90-1/2/3/5 (Brf1) | TFIIIB90-1/2/3/5 (Brf1), a1922 | Santa Cruz Biotech, 200ug/mL | Mouse, Monoclonal, 1:500 |
| RIG-I | RIG-I (D-12), G1822 | Santa Cruz Biotech, 200mg/mL | Mouse, Monoclonal, 1:1000 |
| RIG-I | RIG-I (D14G6), 3743S | Cell Signaling Technology, 71ug/mL | Rabbit, Monoclonal, 1:1000 |
| RIG-I | RIG-I/DDX58, 20566-I-AP | ProteinTech, 1000ug/mL | Rabbit, Polyclonal, 1:1000 |
| MAVS | MAVS (E827M), 83000S | Cell Signaling Technology, 75 ug/mL | Rabbit, Monoclonal, 1:1000 |
| GAPDH | GAPDH, 2793304 | Invitrogen, 100ug/mL | Mouse, Monoclonal, 1:3000 |
| Goat Anti-Mouse IgG | Goat Anti-Mouse IgG (H+L) Human ads-HRP, 1031-05 | Southern Biotech | Goat, Polyclonal, 1:5000 |
| Goat Anti-Rabbit IgG | Goat Anti-Rabbit IgG-HRP, 4030-05 | Southern Biotech | Goat, Polyclonal, 1:5000 |

### RT-qPCR

Total RNA was isolated from cells using TRIzol (Invitrogen 15596018) and treated with Turbo DNase (Invitrogen AM2239) to remove genomic DNA contamination. DNase-treated RNA was reverse transcribed using SuperScript III (SSIII) (Thermo Fisher 18080044) with random 9-mer primers (IDT) for total RNA quantification. Gene expression was assessed by amplifying cDNA with iTaq SYBR Green Supermix (Bio-Rad 1725125) on a QuantStudio 3 instrument (Applied Biosystems A28566). Primer sequences are provided in Table 1. Quantification was performed using the delta delta CT (ΔΔCt) method, and average fold changes were calculated relative to the 18S internal control.

### MHV68 viral titration

Viral supernatants were collected at the indicated time points (0-, 1-, 2-, and 3-days post-infection, dpi) and stored at-80°C until analysis. For all titration assays, 5.0 x 10^3^ NIH 3T3 cells/well were seeded into 96-well culture plates 24 h prior to infection.

Supernatants were pre-diluted at 1:1, 1:100, or 1:1000 depending on the dpi and expected viral load. Using non-racked cluster 12-tube strips (Research Products International 162529) supernatants were serially diluted 1:5 in DMEM + 10% FBS. Serial dilutions were transferred onto cells and incubated for 7 days at 37°C. At 7 dpi, wells were scored for cytopathic effect (CPE) and GFP positivity. FFU/mL and TCID50/mL were calculated using the Spearman and Kärber method.

### Plaque size assay

NIH3T3 cells were seeded in 6-well plates at a density of 5.0 x 10^5^ cells/well. 24 hours after seeding, cells were nucleofected with either non-targeting or Brf1-targeting siRNA at a final concentration of 200 nM per well. 24 hours post-nucleofection, cells were infected with MHV68-GFP at an MOI of 0.0025 for 90 minutes. Following initial infection, the inoculum was removed, and cells were washed twice with 1X PBS before overlaying with 2 mL of 1% methylcellulose in DMEM. At 3 days post-infection, total plaques were counted to assess viral entry. For plaque size analysis, 25 plaques per condition were imaged. Image files were randomized prior to analysis, and plaque areas were quantified using pixel area with FIJI software.

### Extracellular genome copy assay

Viral supernatants harvested from corresponding experiments were subjected to DNase treatment at 37°C for 30 minutes. DNase treatment was then inactivated with 50mM EDTA at 65°C for 10 minutes. Virion capsids in the supernatant were then lysed using 400 µg/mL proteinase K and 10% SDS incubated at 37°C for 2 hours with constant agitation. Phenol chloroform extraction was then performed on the lysed virions to extract viral genomic DNA. Following extraction gDNA was purified using ethanol precipitation and resuspended in 20 µl of sterile water. Genome copies were then quantified by qPCR using gB (ORF8) primers. A standard curve was generated using MHV68 BAC 168 plasmid with known genome equivalents to calculate extracellular viral genome copies for each sample.

### Flow Cytometry

Cells were collected and incubated in 4% PFA for 15 minutes for fixation. Cells were washed twice in PBS and then resuspended in FACS buffer for flow cytometry analysis. Flow Cytometry was performed on Beckman Coulter CytoFLEX. Cells for GFP percentage analysis were gated first by FSC-A vs SSC-A gating for live cells. Singlets were gated using FSC-A vs FSC-H. Singlets were double gated using SSC-A vs SSC-H. GFP positive cells were then gated using FSC-A vs B525A. Cell viability was measured using Zombie Violet (Biolegend) dye as previously described^29^.

### RNA sequencing and transcriptome analysis

Total RNA was submitted to Azenta Life Sciences (GENEWIZ, South Plainfield, NJ, USA) for library preparation, quality control, and sequencing. Libraries were prepared according to the provider’s standard RNA-seq protocols and sequenced on an Illumina platform to generate paired-end reads. Raw FASTQ files were returned for downstream bioinformatic analysis. RNA-seq data were processed using HAROLD (https://github.com/dremellab/HAROLD), a Snakemake^30^ based RNA-seq analysis pipeline. All pipeline steps were executed in versioned Docker/Singularity containers to ensure reproducibility^31^. Raw paired-end reads (150 bp) were first validated with fastQValidator version 0.1.1a to detect truncated or malformed FASTQ files, then adapter-and quality-trimmed with Cutadapt v4.9^32^ (minimum length 15 bp, quality cutoff Q20). Trimmed reads were aligned to a combined reference comprising the mouse genome, MHV68 genome, and ERCC spike-in sequences, using STAR v2.7.6a^33^ in 2-pass mode (--twopassMode Basic; --outFilterMultimapNmax 20 -- outFilterMismatchNmax 2 --outFilterMismatchNoverLmax 0.3 --alignSJoverhangMin 15--alignSJDBoverhangMin 15). Gene-level read counts were obtained directly from STAR’s --quantMode GeneCounts output. Alignments were sorted and indexed with Samtools v1.21^34^, and PCR duplicates were marked with Picard MarkDuplicates v2.27.5. ERCC reads were used to generate standard curves similar to Schertzer, M. D., et al^35^, using their known relative concentrations. All biological replicates had ERCC derived standard curves with R2>0.9. ERCC normalized gene counts were calculated as follows:

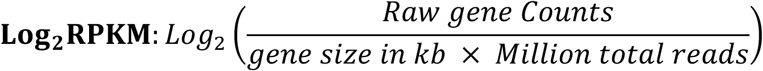

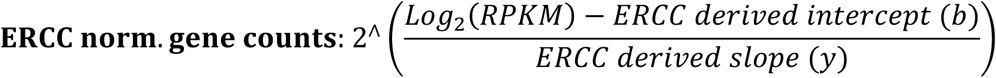

Differentially expressed genes (DEGs) were identified using limma-voom v3.62.2^36^. ERCC normalized counts were transformed using the voom function to estimate mean-variance weights, followed by linear modeling (lmFit) according to the specified group comparison and empirical Bayes moderation (eBayes) to assess differential expression. Genes were classified as differentially expressed using a log2 fold-change threshold of 1.0 and p-value threshold of 0.05 (Benjamini-Hochberg correction). Volcano plots were limited to mouse protein-coding genes. Pathway analysis was performed using pre-ranked gene set enrichment analysis (GSEA) via the clusterProfiler:GSEA function (fgsea-based), with genes ranked by their limma-voom differential expression statistic and tested against the MSigDB Hallmark (H) gene set collection using msigdbr. Gene sets were filtered to a minimum size of 15 and maximum size of 500, with enrichment considered significant at a nominal p-value cutoff of 0.05.

### Genome Assemblies

Mouse: mm39, gencode.vM29^37^

ERCC Spike-In: available from ThermoFisher (#4456740)

MHV68: MH636806.1^38^ modified to remove the beta-lactamase gene (Δ103,908-105,091), CDS annotation used for transcript quantification. MHV68 transcripts were clustered into the following temporal classes: Immediate Early (IE)-ORF50; Early (E)-M4, ORF6, ORF10, ORF30, ORF36-37, ORF44, ORF48, ORF54, ORF56-57, ORF59-61, ORF75A; Leaky Late (L1)-M1, M3, M5, M6, M14, ORF7, ORF9, ORF11, ORF18, ORF21, ORF24, ORF27, ORF32, ORF34-35, ORF38, ORF40, ORF42, ORF46, ORF58, ORF63, ORF68, ORF72; Late (L)-M7, M9, M15, M17, M18, ORF4, ORF8, ORF17, ORF19-20, ORF22-23, ORF26, ORF29, ORF33, ORF39, ORF43, ORF45, ORF49, ORF52-53, ORF55, ORF62B, ORF64, ORF66-67, ORF69^38, 39^.

### Statistical analysis

All experiments were performed with at least three biological replicates, defined as independent experiments conducted on different days using distinct cell populations (separate stock vials or passage numbers). RT-qPCR data were analyzed using raw ΔCt values. The statistical tests applied for each experiment are specified in the figure legends.

### Data Availability

RNA-seq data has been deposited under BioProject PRJNA1521517. Plasmids generated have been deposited with Addgene. Code for RNA-seq analysis can be found at https://github.com/dremellab/HAROLD.

## RESULTS

### siRNA-mediated Brf1 knockdown is sustained over multiple rounds of MHV68 infection

DNA virus infection has been shown to stimulate transcription of Pol III genes, particularly at those made from Type I (e.g., 5S rRNA) and Type II promoters (tRNAs, B2 SINEs, 7SL, etc.) ^17, 21, 40, 41^. To limit RNA Polymerase III (Pol III) transcription from Type I and II promoters, while still allowing Pol III transcription to occur at other essential genes with Type III promoters (e.g., 7SK and U6), we targeted Brf1, an essential component of the Transcription Factor IIIB (TFIIIB) complex (Fig 1A). NIH 3T3 mouse fibroblasts were transfected with either non-targeting (NT) or a Brf1-specific siRNA pool to decrease Brf1 protein expression. Due to the essentiality of Brf1, we assessed cell viability using Zombie viability dye and flow cytometry each day for four days following transfection (Fig 1B). No significant differences were observed between NT-and Brf1-siRNA-treated cells, indicating that Brf1 knockdown did not affect NIH 3T3 cell viability during the time frame measured. To validate the loss of Brf1 protein with siRNA treatment, we performed western blot analysis on uninfected cells across four days post-transfection (Fig 1C). Brf1-siRNA-treated cells displayed a consistent and robust reduction (∼95%) in Brf1 protein levels relative to NT control cells. We next assessed whether MHV68 infection influenced knockdown efficiency by infecting cells at an MOI of 0.05 at 24 hours post siRNA treatment (Fig 1D). Brf1 protein remained depleted throughout the course of infection, with minimal Brf1 protein expression evident through three days post-infection. These data confirm that siRNA-mediated depletion of Brf1 is stable in both uninfected and infected cells.

**Figure 1.**
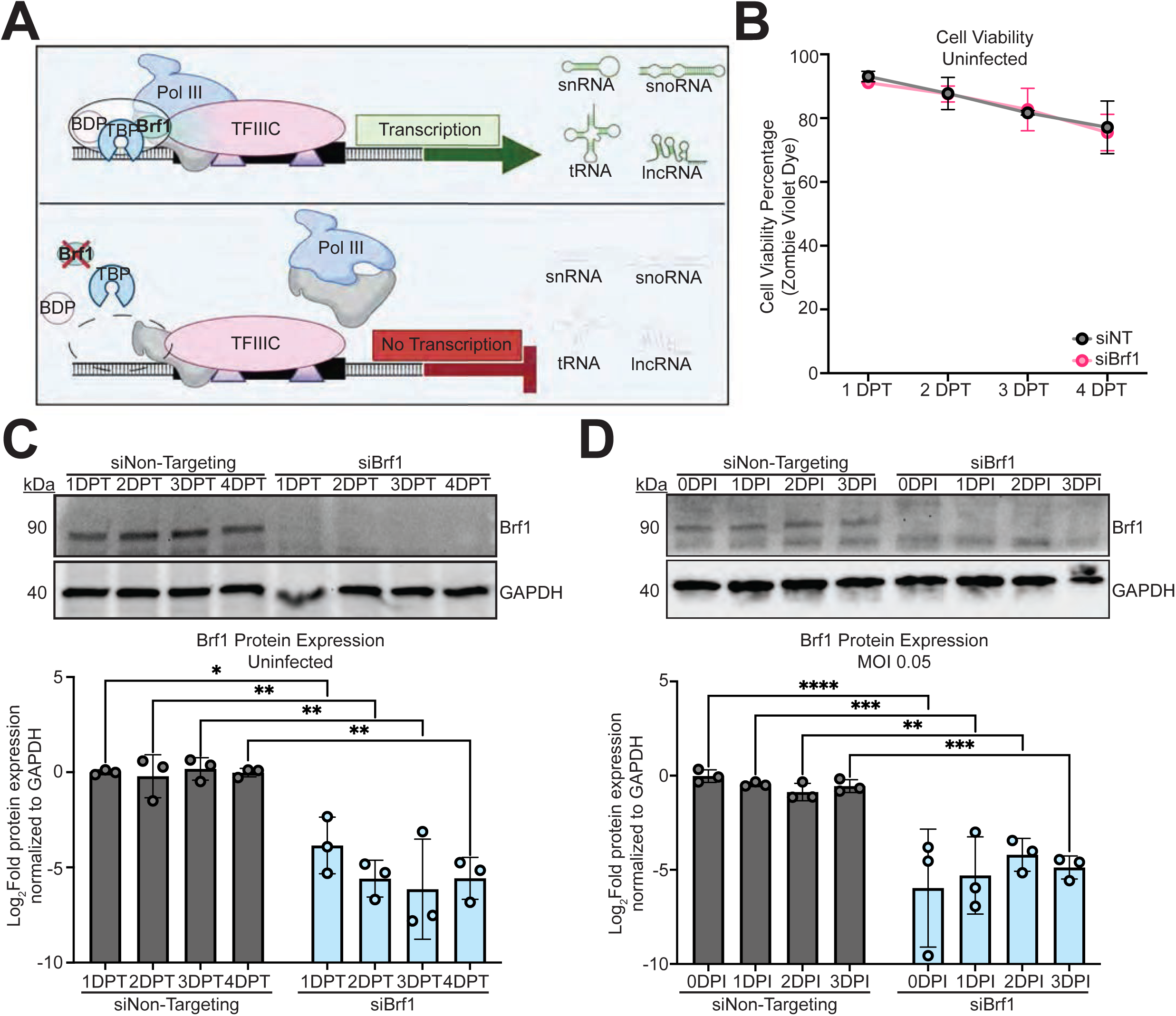
Validation of Brf1 knockdown in NIH 3T3 cells. (A) Schematic illustrating RNA polymerase III (Pol III) transcription under WT conditions (top) and following Brf1-targeting siRNA treatment (bottom). (B) Cell viability of NIH 3T3 cells transfected with non-targeting (siNT) or Brf1-targeting (siBrf1) siRNA, measured by Zombie Violet staining and flow cytometry 1-4 days post-transfection (dpt). Representative western blot and densitometric quantification of Brf1 protein expression in (C) uninfected NIH 3T3 cells following siNT or siBrf1 transfection at 1-4 dpt and (D) infected NIH 3T3 cells (MOI = 0.05) following siNT and siBrf1 transfection at 0-3 days post-infection (dpi). GAPDH was used as a loading control. Data represents three independent experiments and is shown in a log_2_ scale. Statistical significance was determined by one-way ANOVA. P ≤ 0.05 (*), P ≤ 0.01 (**), P ≤ 0.001 (***), P ≤ 0.0001 (****).

### Profiling RNA Pol III promoter-dependent effects of Brf1 knockdown during MHV68 infection

To confirm that Brf1 knockdown limits Pol III transcription, we quantified representative Pol III transcripts from each promoter type following siRNA treatment and MHV68 infection at an MOI 0.05 (Fig 2). As mentioned previously, both Type I and II promoters rely on Brf1-dependent formation of the TFIIIB complex for Pol III recruitment. Among Type I and II Brf1-dependent transcripts, levels of premature tRNAs, tyrosine and leucine, were significantly reduced in Brf1-depleted cells, while 5S rRNA and 7SL RNA levels remained unchanged (Fig 2). We reasoned that the abundance and stability of pre-existing pools of 5S rRNA and 7SL obscure any decrease in nascent transcripts following Brf1-depletion. In support of this, the Ct values of steady state 5S and 7SL show they are approximately 10,000-fold more abundant than pre-tRNAs, which likely results in minimal loss of signal upon Brf1 knockdown, at least over the time course measured (Fig S1A). In contrast, Type III promoters are Brf1-independent, recruiting TFIIIB assembly with Brf2 instead of Brf1. Compared to Type I and II transcripts, Type III transcripts showed a modest increase in Brf1-depleted cells, but only at 3 dpi with MHV68, indicating a synergistic effect of Brf1 knockdown and infection on Type III transcript expression (Fig 2). Together, these results indicate that decreased Pol III activity following our siRNA treatment is specific and limited to Brf1-dependent genes.

**Figure 2.**
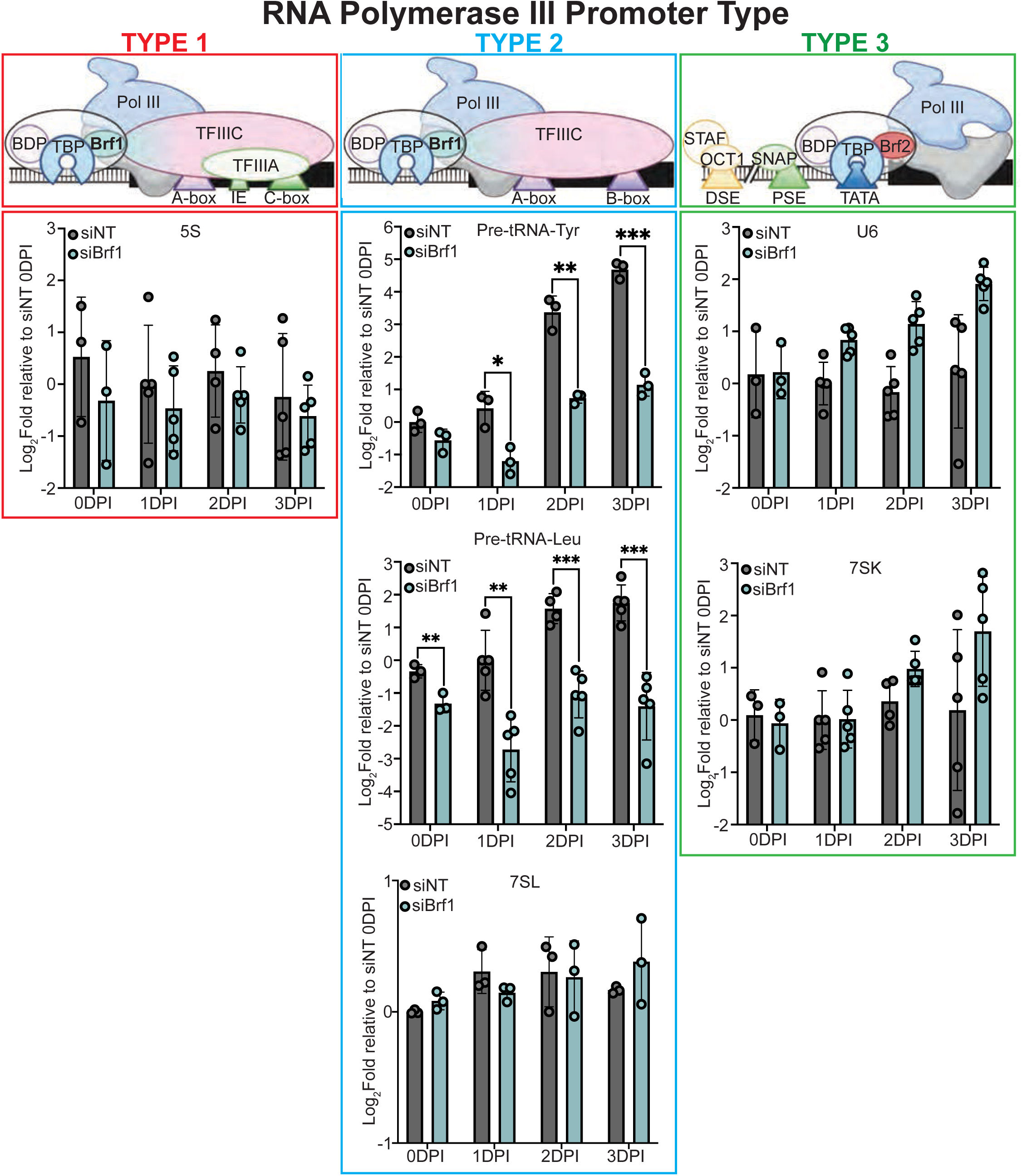
Non-coding RNA expression during MHV68 infection following Brf1 knockdown. Top portion shows schematics of protein complexes required for RNA Pol III recruitment separated by promoter type. Bottom portion shows RT-qPCR analysis of RNA polymerase III-transcribed non-coding RNAs in NIH 3T3 cells transfected with non-targeting (siNT) or Brf1-targeting (siBrf1) siRNA during MHV68 infection (MOI = 0.05). Transcripts are groups by promoter type: Type 1 (5S rRNA; red), Type 2 (Pre-tRNA-Tyr, Pre-tRNA-Leu, and 7SL RNA; blue), and Type 3 (U6 and 7SK RNA; green). Samples were collected from 0-3 days post-infection (dpi). Data represent 3-5 independent experiments. Log2 fold change was calculated using the siNT 0 dpi sample as the relative sample. Statistical significance was determined by one-way ANOVA. P ≤ 0.05 (*), P ≤ 0.01 (**), P ≤ 0.001 (***), P ≤ 0.0001 (****)

### Brf1-dependent Pol III activity limits viral transcription and protein production during MHV68 infection

Having established that Brf1 dependent Poll III activity is limited during MHV68 infection, we next evaluated how Brf1 depletion affects viral transcription and protein production during a low-MOI (0.05) infection. Viral gene expression was assessed by RT-qPCR using the MHV68 early gene ORF50, which encodes the replication and transcriptional activator (RTA), and the late gene gB, which encodes a conserved herpesvirus glycoprotein (Fig 3A). Brf1-depleted cells exhibited significantly increased ORF50 transcript levels (ranging from 2-5-fold higher) at 1-and 2-days post-infection (dpi) compared to cells treated with non-targeting siRNA, with a modest but more variable increase observed at the endpoint of infection (3 dpi). gB levels were similarly increased at 1-and 2-days post infection (ranging from 1-5-fold higher) in Brf1-depleted cells but were reduced (1.5-fold) relative to control infection at 3 days post-infection. These data indicate that Brf1 depletion enhances viral transcript levels during the early and mid-stages of a low MOI/multi-step MHV68 infection. To look at total changes in viral RNA abundance, RNA sequencing was performed on the time course infection to assess all MHV68 transcripts (Fig 3B; Fig S2A). Fold change values for ORF50 and gB from RNA-seq show an increase (ranging from 2-4-fold higher) at 1-and 2-days post-infection in Brf1-deficient cells, similar to our RT-qPCR data (Fig 3B; Fig S2B).

**Figure 3.**
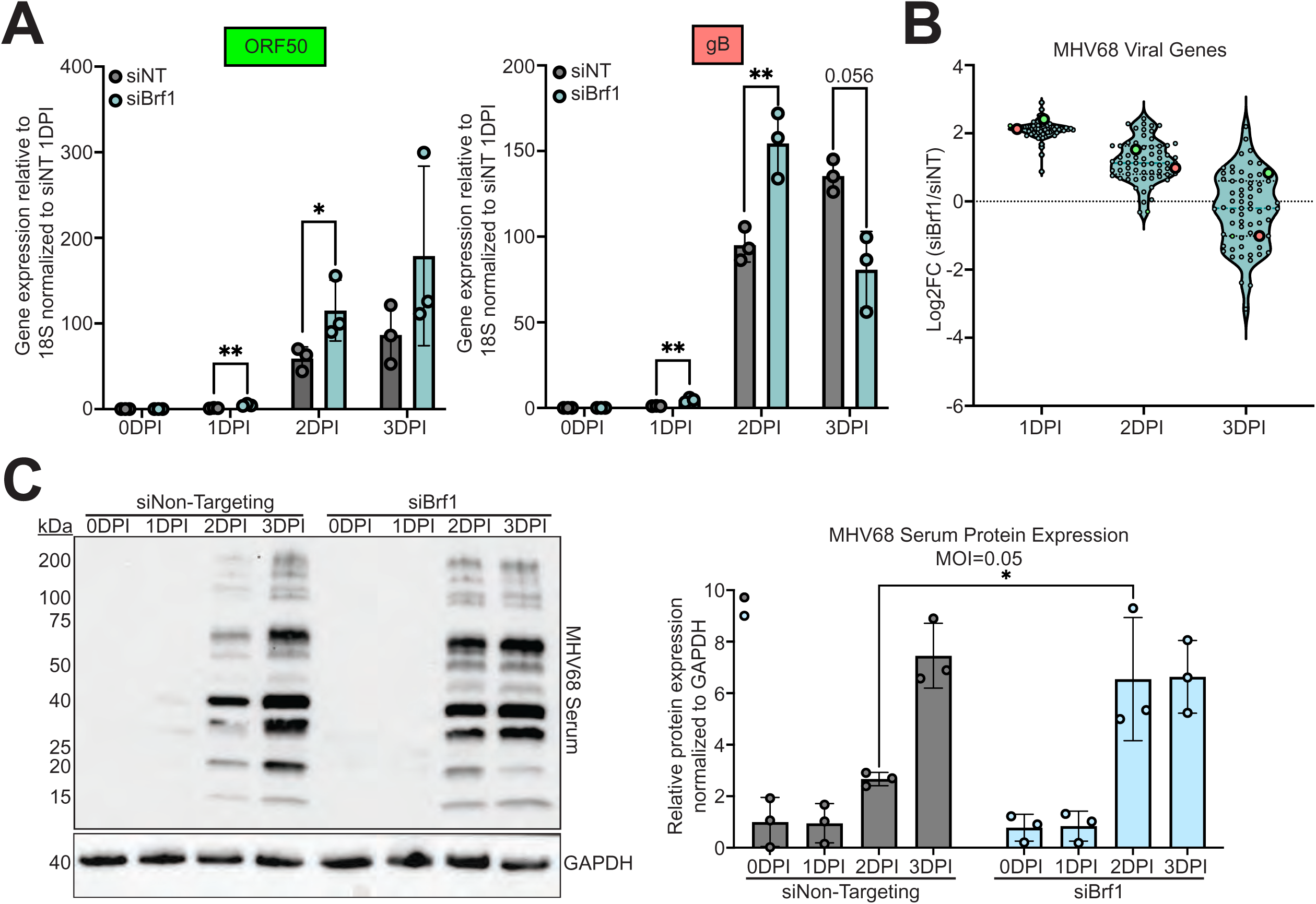
Brf1 knockdown alters MHV68 viral gene expression and protein accumulation. NIH 3T3 cells were transfected with non-targeting (siNT) or Brf1-targeting (siBrf1) siRNA and infected with MHV68-GFP (MOI=0.05). (A) RT-qPCR analysis of viral gene expression from RNA collected at 0-3 days post-infection (dpi). Expression levels of the early viral gene ORF50 (green) and the late viral gene gB (red) were quantified. (B) Log_2_ fold change of normalized transcript abundance determined by bulk RNA sequencing from the same RNA samples collected at 0-3 dpi. Green and red symbols represent ORF50 and gB, respectively. (C) Representative Western blot and corresponding densitometric quantification of total MHV68 viral protein accumulation from 0-3 dpi. GAPDH was used as a loading control. Statistical significance was determined by one-way ANOVA. P ≤ 0.05 (*), P ≤ 0.01 (**), P ≤ 0.001 (***), P ≤ 0.0001 (****)

To determine whether increased transcript levels corresponded to changes in viral protein levels, we analyzed total viral protein using a polyclonal anti-MHV68 serum (Fig 3C). Minimal viral protein was detected at 0-and 1-days post-infection in both conditions; however, a significant increase in total viral protein (2.7-fold) was observed in Brf1-depleted cells at 2 days post-infection. By the endpoint of infection, viral protein largely stabilized, with only modest differences seen between siRNA treatments. These results parallel the transcriptional dynamics, supporting a model in which the knockdown of Brf1 dependent Pol III activity promotes viral gene expression and protein production over the course of MHV68 infection, with effects diminishing approaching the endpoint of infection.

### Loss of Brf1-dependent Pol III activity leads to increased MHV68 infection at a low MOI

To directly assess how reduced Pol III activity influences MHV68 replication, we quantified virion production by Tissue Culture Infectious Dose 50% (TCID_50_) assays with supernatant collected at multiple timepoints post-infection. Viral titers were comparable between control and Brf1-depleted cells at 0-and 1-dpi (Fig 4A). In contrast, a significant divergence emerged at later time points: at both 2 and 3 dpi, Brf1-depleted cells exhibited a 5-10-fold increase in viral titers relative to control infection (Fig 4B).

**Figure 4.**
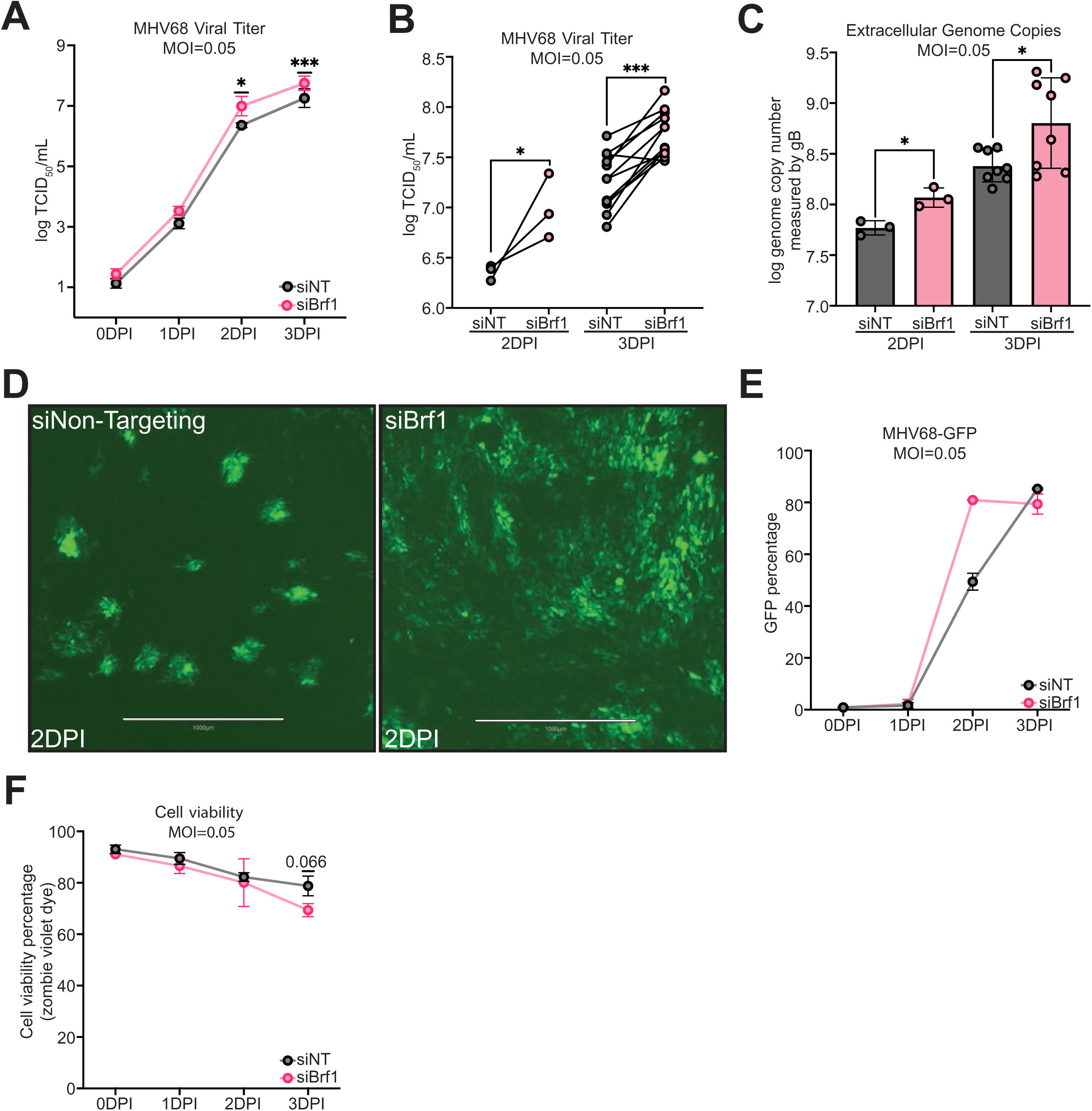
Brf1 knockdown enhances MHV68 infection output in NIH 3T3 cells. NIH 3T3 cells were transfected with non-targeting (siNT) or Brf1-targeting (siBrf1) siRNA and infected with MHV68-GFP (MOI=0.05). (A) Viral titers measured by TCID_50_ assay from 0 to 3 days post-infection (dpi). (B) Individual paired biological replicates from the 2-and 3-dpi time points shown in (A). (C) Extracellular genome copies quantified by qPCR measured gB using genomic DNA isolated from harvested culture supernatants at 2 and 3 dpi. (D) Representative fluorescence images of MHV68-GFP infection at 2 dpi in siNT-and siBrf1-treated cells. (E) Flow cytometry measuring GFP percentage arising from MHV68-GFP infection from 0 to 3 dpi. N=2 (F) Cell viability from 0 to 3 dpi, measured by Zombie Violet staining and flow cytometry. Statistical significance was determined by unpaired T-test and one-way ANOVA. P ≤ 0.05 (*), P ≤ 0.01 (**), P ≤ 0.001 (***), P ≤ 0.0001 (****)

Extracellular genome copies were measured by qPCR at 2-and 3-dpi using primers specific for viral gene, gB. Modestly increased genome copies were observed (2-fold) at 2-days post infection and significantly (3.8-fold) at 3-days post infection in Brf1-deficient cells, corresponding to increased viral titers seen from TCID_50_ assays (Fig 4C). This increase in infectious virion production corresponded with distinct infection phenotypes visualized in MHV68-GFP-infected cultures at 2 dpi (Fig 4D). Control cultures displayed dense but localized foci of infection, whereas Brf1-depleted cultures showed a more dispersed pattern of infection, while maintaining robust GFP intensity. Further, flow cytometry analysis of GFP% cells indicate an almost 2-fold increase in GFP-positive cells when comparing Brf1-deficient to control cells at 2 dpi (Fig 4E). Assessment of cell viability at the endpoint of infection revealed a modest increase in cell death in Brf1-depleted cells, consistent with enhanced viral replication and cytopathic effects (Fig 4F). Together, these results indicate an anti-viral role for Brf1 dependent RNA polymerase III activity during MHV68 infection.

### siRNA resistant Brf1 blocks enhancement of MHV68 infection regardless of siRNA treatment

To confirm that the observed antiviral phenotype is specifically attributable to Brf1 function rather than siRNA off-targeting effects, we employed a genetic rescue approach (Fig 5). NIH3T3 cells were transduced with lentivirus encoding either an empty control vector or a codon-optimized and siRNA-resistant Brf1 overexpression construct (Brf1-CO). Successful overexpression was confirmed by western blot analysis (Fig 5A). Brf1-CO cells showed no difference in pre-tRNA-Tyr expression between non-targeting and Brf1-targeting siRNA conditions, confirming that exogenous Brf1 is functional and capable of restoring Pol III–dependent transcription (Fig 5B). We observed a modest decrease in Brf1-CO levels with Brf1-targeting siRNAs, suggesting that codon optimization did not fully block siRNA silencing. However, Brf1-CO expression in the presence of siRNA remained far above native Brf1 levels and was capable of fully rescuing pre-tRNA expression (Fig 5A-B). Brf1-CO cells showed no significant difference in viral protein accumulation, viral titers, or viral gDNA copies between non-targeting and Brf1-targeting siRNA treatment, indicating that the antiviral phenotype is specific to Brf1 expression (Fig 5C, D, E). We also note that viral protein accumulation and viral titers in Brf1-CO cells were increased relative to empty vector control cells under both siRNA conditions, suggesting that elevated Brf1 expression may enhance viral replication independent of siRNA targeting (see discussion).

**Figure 5.**
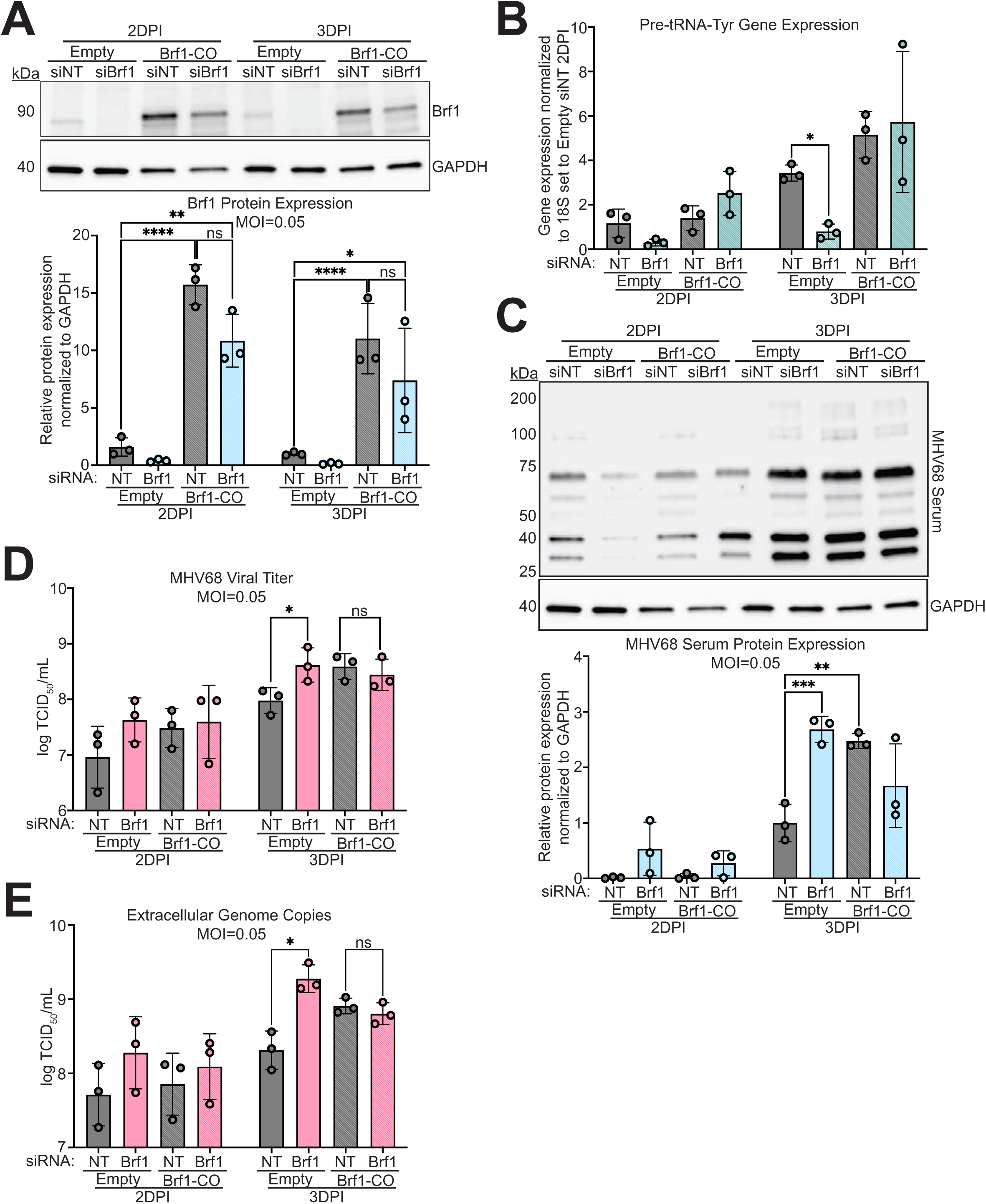
Rescue of phenotype using an siRNA-resistant Brf1 expression construct. NIH 3T3 cell lines stably transduced with either an empty vector control or a codon-optimized, siRNA resistant Brf1 expression construct were transfected with non-targeting (siNT) or Brf1-targeting (siBrf1) siRNA and infected with MHV68-GFP (MOI=0.05). Cells and supernatants were harvested at 2-and 3-days post-infection (dpi) for downstream analyses. (A) Representative western blot and densitometric quantification of Brf1 protein levels, shown in log2 scale. (B) RT-qPCR analysis of pre-tRNA-Tyr expression. (C) Representative western blot and densitometric quantification of total MHV68 viral protein accumulation. (D) Viral titers determined by TCID_50_ assay. (E) Extracellular viral genome copies quantified by qPCR using genomic DNA isolated from harvested supernatants. GAPDH used as loading control for all western blot analyses. Statistical significance was determined by one-way ANOVA. P ≤ 0.05 (*), P ≤ 0.01 (**), P ≤ 0.001 (***), P ≤ 0.0001 (****)

Together, these findings indicate that reduced Pol III activity-achieved through Brf1 depletion promotes enhanced MHV68 replication during a low MOI infection, aligning with observed increases in viral gene expression and protein accumulation, and is not due to off-targeting effects of siRNA treatment.

### Overexpression of select Brf1-dependent RNA Pol III transcripts does not alter MHV68 titer

Because Brf1-dependent RNA Pol III activity appears to restrict MHV68 replication, we next hypothesized that a specific Pol III transcript might mediate this antiviral phenotype. Overexpression vectors were generated for transcripts made in a Brf1-dependent manner, including 5S (Type I promoter), pre-tRNA-Arg, pre-tRNA-Tyr, B2 SINE, and 7SL (all Type II promoters). We also constructed a 7SK overexpression vector (Type III promoter, not dependent on Brf1) as a control. Each transcript is expressed through an external modified H1/7SK hybrid M11 promoter to boost Pol III transcription^43^ (Fig S1B). While we could detect a significant increase in pre-tRNAs and 7SK transcript levels, this was not achievable for 5S, B2 SINE, and 7SL, likely due to background abundance of these non-coding RNAs (Fig S1C). Following transfection with the overexpression vectors and a low MOI MHV68 infection, no significant change in MHV68 titer was observed. We did observe a trending increase in viral titer with 7SK overexpression. In contrast, extracellular genome copies with 7SK overexpression were decreased, resulting in a lower particle/PFU ratio (higher infectivity of virus particles) when 7SK is overexpressed (Fig S1F). To assess the role of 7SK further, we used antisense oligonucleotides (ASOs) previously described^44^ to limit transcript levels to determine how MHV68 infection is affected. Two ASOs were used, one targeting the 5’ end and one targeting the 3’ end of 7SK transcripts, for degradation by RNase H1 (Fig S1G). 7SK transcript levels were significantly decreased by both ASOs compared to scramble control as early as 5 hours post-transfection and remained down until 3 days post-infection (Fig S1H). While 7SK transcript were down, we saw no significant difference in MHV68 viral titers, but a modest increase in extracellular genome copies (Fig S1I, J). This resulted in a significantly higher particle/PFU ratio (Fig S1K) (lower infectivity of virus particles). These data together show that 7SK may play an important pro-viral role in the system (see discussion).

### Brf1 knockdown minimally impacts MHV68 replication following inoculation at a high MOI

To determine whether the increases in viral titers, gene expression, and protein accumulation seen during multiple rounds of infection were driven by enhanced viral replication, we infected Brf1-deficient cells with MHV68 at a high MOI (5) (Fig 6). Given the 18–24-hour lytic cycle of MHV68^45^, samples were collected at multiple timepoints throughout infection up to 24 hours post-infection (hpi). Western blotting for Brf1 protein and qPCR analysis of pre-tRNA-Tyr confirmed efficient Brf1 knockdown and the expected loss of Pol III activity (Fig 6A, B).

**Figure 6.**
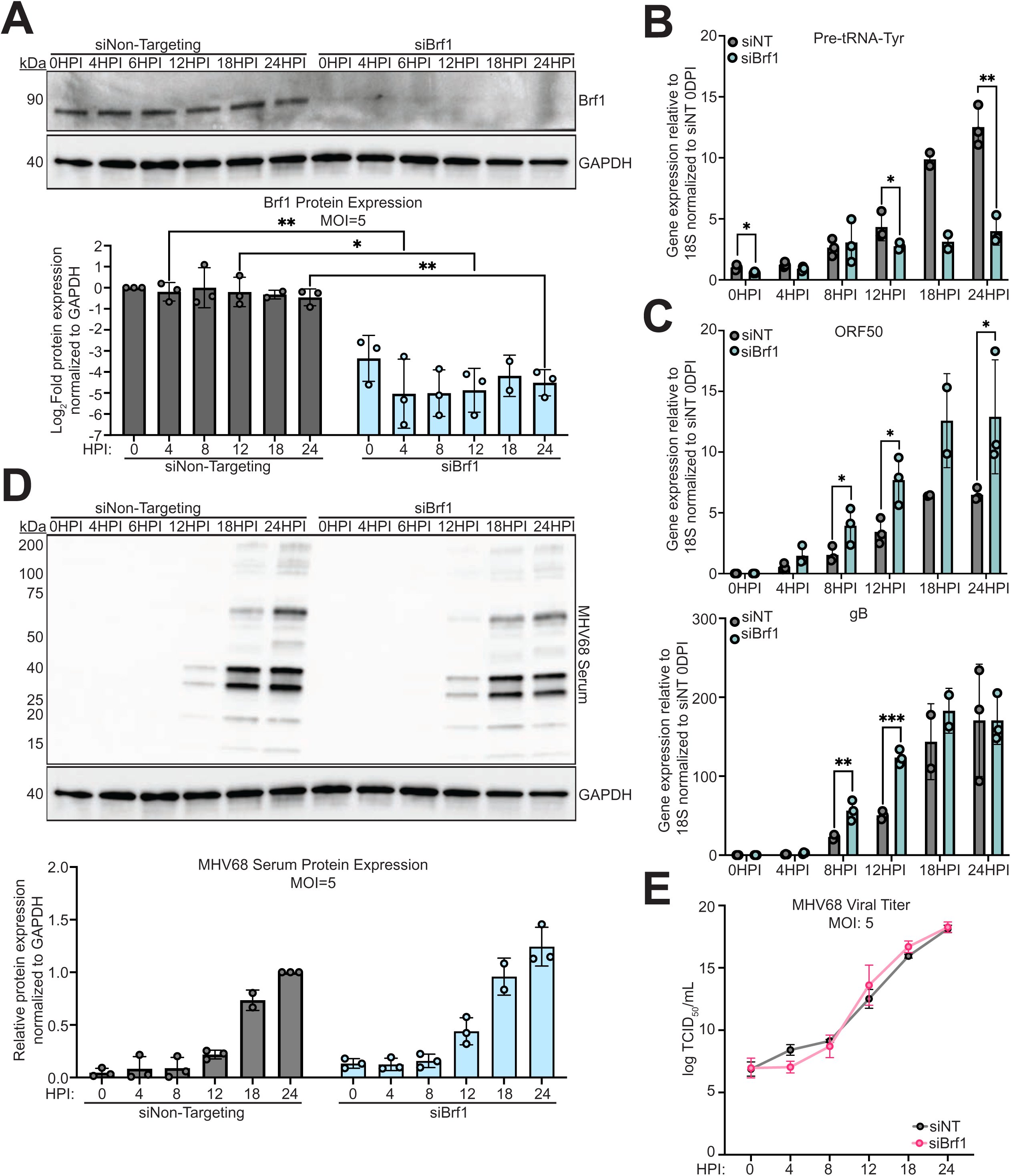
Brf1 knockdown does not alter a high-MOI MHV68 infection. NIH 3T3 cells were transfected with non-targeting (siNT) or Brf1-targeting (siBrf1) siRNA and infected with MHV68-GFP (MOI = 5). Cells were harvested at indicated time points up to 24 hours post-infection (hpi). (A) Representative western blot and densitometric quantification of Brf1 protein levels. (B) RT-qPCR analysis of pre-tRNA-Tyr gene expression. (C) RT-qPCR analysis of the viral early gene, ORF50, and the late gene, gB. (D) Representative western blot and densitometric quantification of total MHV68 viral protein accumulation. (E) Viral titers determined by TCID_50_ assay. GAPDH served as a loading control for all western blot analyses. Statistical significance was determined by one-way ANOVA. P ≤ 0.05 (*), P ≤ 0.01 (**), P ≤ 0.001 (***), P ≤ 0.0001 (****)

Under high-MOI conditions, early viral gene expression (ORF50) was consistently elevated in Brf1-deficient cells beginning at 8 hpi and continuing to the endpoint of infection. Late viral gene expression (gB) was also increased at 8 and 12 hpi, but these converged with control cells by the later stages of infection (Fig 6C). Viral protein accumulation became detectable at 12 hpi and stabilized at 18 hpi; however, no substantial differences in protein abundance was observed between control and Brf1-deficient cells at any time point (Fig 6D). Assessment of infectious virion production by TCID50 revealed only a modest increase in viral titers at 12 hpi in Brf1-deficient cells, which normalized by 18 and 24 hpi (Fig E). These findings indicate that the enhanced viral titers observed under low-MOI conditions are only minimally recapitulated during a high-MOI infection, suggesting that functional role of Brf1-dependent Pol III activity is most evident during a low-MOI infection.

### RNA sequencing reveals accelerated interferon induction followed by enhanced host shutoff in Brf1-deficient cells during infection

To identify transcriptional mechanisms underlying the enhanced viral phenotypes observed in Brf1-deficient cells, RNA collected from MHV68 time course experiments was subjected to bulk RNA sequencing. Differential expression analysis was performed by comparing Brf1-deficient cells to control cells at 0-3 days post infection (dpi) using log_2_ fold change of normalized read counts (Fig 7). At 0 dpi, which corresponds to 24 h post siRNA treatment, minimal differences in transcript abundance were observed between both conditions, indicating that Brf1 depletion alone at this timepoint had limited impact on basal gene expression. By contrast, at 1 dpi, substantial transcriptional changes emerged with over 800 genes being dysregulated between the two conditions. At later time points (2-and 3 dpi) a global reduction in host mRNA abundance was observed, consistent with host shutoff ^46^ (Fig 7A). Notably, this decrease in transcript levels appears more pronounced in Brf1-deficient cells. Total read percentage was measured for both control and Brf1 deficient cells. While host genes accounted for the majority of reads at 2 dpi in controls cells, we saw a significant shift in Brf deficient cells with about 70% of total reads being viral (Fig 7B). This is consistent with the increase in MHV68-infected cells in these cultures, as measured by GFP-positivity (Fig 4E).

**Figure 7.**
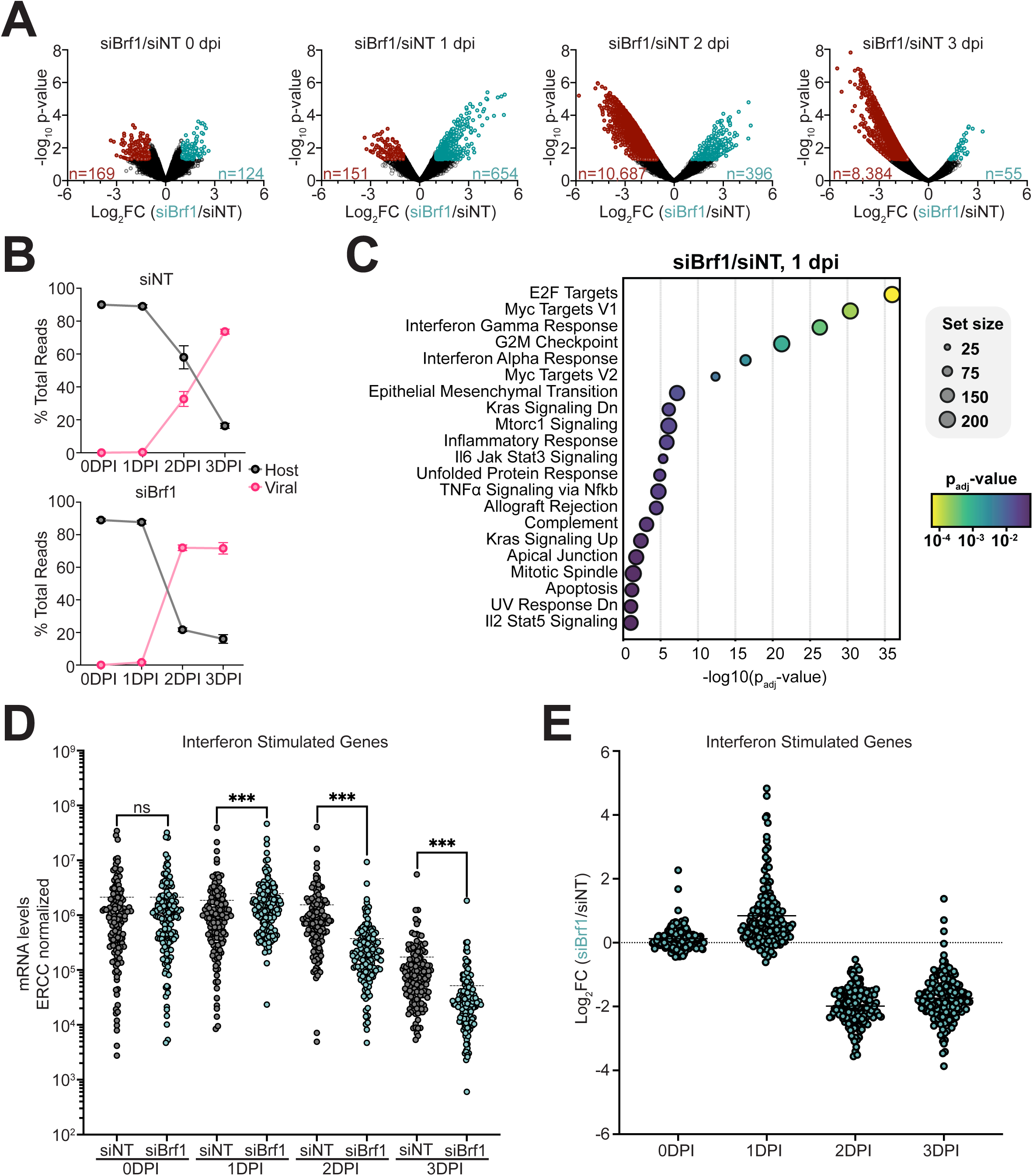
RNA sequencing analysis shows enhanced viral host shutoff and faster induction of interferon response. Bulk RNA sequencing performed on 3 biological replicates. (A) Log2 fold change analysis of host genes measuring Brf1 deficient cells over control. Each graph represents timepoints 0-3 days post infection. (B) Total read percentage of host and viral mRNA at 0-3 dpi. Non-targeting siRNA-treated cells and Brf1-deficient cells are shown on separate graphs. (C) Pathway analysis of 1 dpi showing highly dysregulated pathways based on P_adj_ value (color) and set size (size of circle). (D) ERCC normalized mRNA levels of interferon related genes upregulated at 1 dpi in Brf1 deficient cells. (E) Log2 fold change of interferon related genes comparing Brf1-deficient cells to control cells. Statistical significance was determined by one-way ANOVA. P ≤ 0.05 (*), P ≤ 0.01 (**), P ≤ 0.001 (***), P ≤ 0.0001 (****)

To further characterize transcriptional differences at early stages of infection, genes significantly upregulated at 1 dpi in Brf1-deficient cells were analyzed. Interestingly, more than 80 dysregulated genes (Fig S2C) were associated with interferon signaling pathways, including both canonical interferon pathway components and interferon-stimulated genes (Fig 7C). Top hits also included E2F and Myc pathways that are known to be hijacked during gammaherpesvirus infection to induce cell survival and proliferation^47–49^. Myc has also been shown to regulate RNA Pol III through binding of TFIIIB^50^. To directly assess the kinetics of interferon pathway activation, normalized read counts at each time point were compared respective to the 0 dpi baseline within each condition (Fig 7D). At 1 dpi, control cells showed minimal induction of interferon-responsive genes relative to baseline. In contrast, Brf1-deficient cells displayed robust upregulation of these genes at 1 dpi, suggesting a faster and/or stronger interferon response in this culture. To validate these findings, RT-qPCR was performed for two interferon-inducible GTPases, Irgm1 and Irgm2 (Fig S2E). In both cases, gene expression increased during infection; however, induction was detectable at 1 dpi in Brf1-deficient cells, whereas in control cells, upregulation was delayed until 2-dpi. These results support an earlier induction of interferon-responsive genes in Brf1-deficient cells during MHV68 infection. However, by days 2-3 post infection, interferon-responsive gene expression is lower in siBrf1-treated cells, likely supporting the overall higher titers measured in Brf1-deficient cells (Fig 7D-E).

Host pol III transcripts have been shown to trigger MAVS-dependent pathways during herpesvirus infections through sensing by RIG-I^21, 22^. Brf1 depletion would be expected to decrease expression of the ncRNAs known to bind and trigger RIG-I, or perhaps indirectly alter their maturation. We reasoned that an alteration in host Pol III transcripts could influence the innate immune response and subsequent replication outcomes during infection. To test whether RIG-I/MAVS are important for Brf1 restriction of MHV68, we made RIG-I and MAVS knockout cells (Fig 8). Successful knockout of each protein was confirmed by western blot analysis, which demonstrated complete loss of RIG-I and MAVS expression in their respective cell lines (Fig 8A). To verify functional disruption of the pathway, cells were stimulated with poly(I:C), a viral double-stranded RNA mimic, for 24 hours. As expected, poly(I:C) treatment induced robust expression of the interferon-stimulated genes (ISGs) ISG15 and OAS2 in safe-targeting control cells. In contrast, induction of both ISGs was reduced by more than 90% in RIG-I-and MAVS-deficient cells, confirming loss of functional MAVS-dependent interferon signaling (Fig 8B).

**Figure 8.**
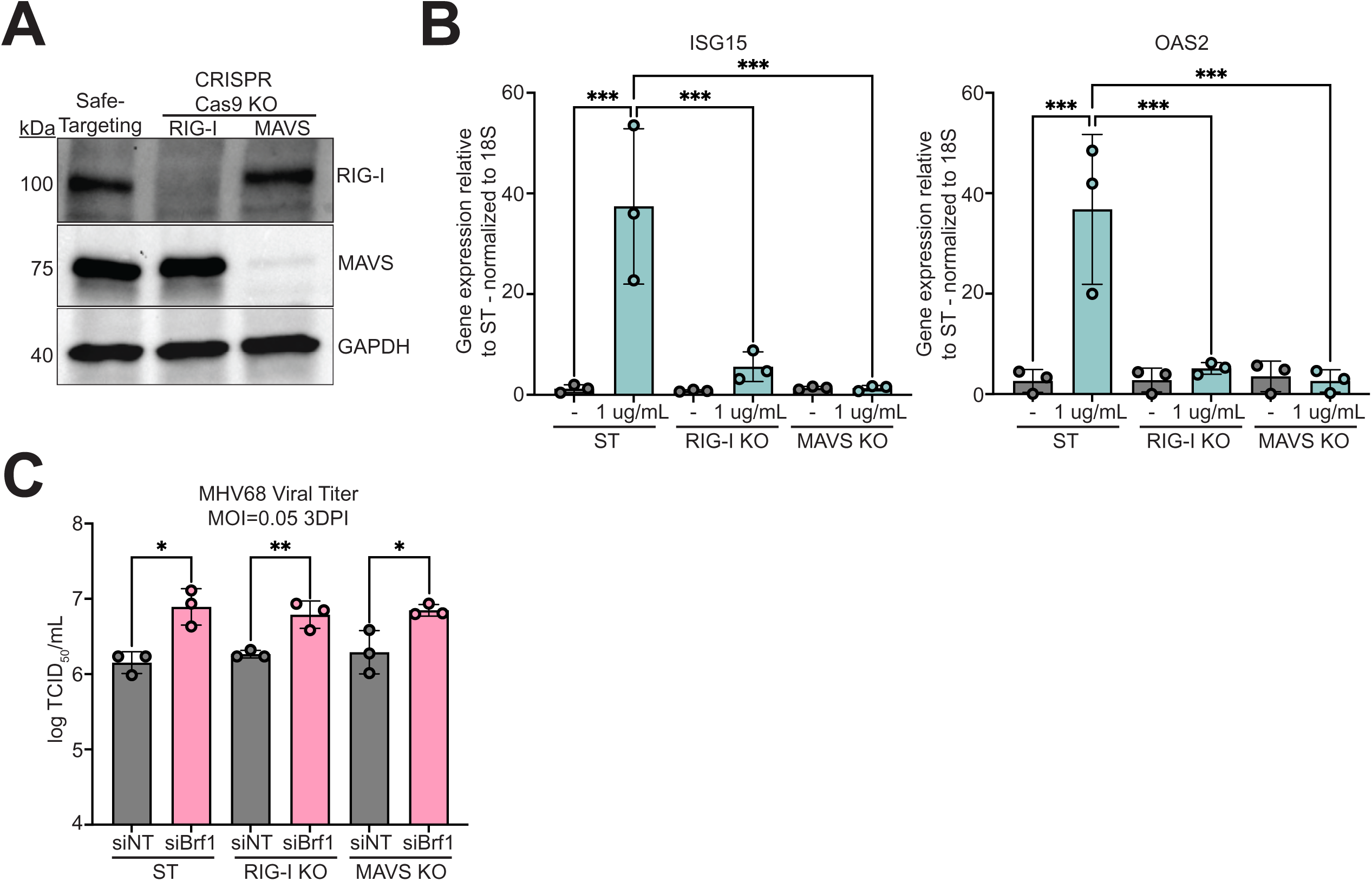
Analysis of the MAVS signaling pathway using RIG-I and MAVS CRISPR-Cas9 knockout cell lines. (A) Representative western blot analysis of safe-targeting control, RIG-I knockout, and MAVS knockout cell lines confirming the loss of target protein expression following CRISPR-Cas9-mediated knockout. (B) RT-qPCR analysis of interferon-stimulated gene (ISG) following stimulation with the dsRNA mimic, poly(I:C). Expression of ISG15 and OAS2 was measured to validate loss of functional MAVS-dependent interferon signaling. (C) Viral titers at 3 days post-infection (dpi) determined by TCID_50_ assay for each indicated cell line. Statistical significance was determined by one-way ANOVA. P ≤ 0.05 (*), P ≤ 0.01 (**), P ≤ 0.001 (***), P ≤ 0.0001 (****)

To determine whether RIG-I or MAVS mediate the effect of Brf1 depletion on viral replication, RIG-I and MAVS knockout cells were transfected with siNT and siBrf1 and subsequently infected with MHV68-GFP at a low MOI. Viral titers were quantified 3 days post-infection using supernatant collected from each condition. Despite the loss of MAVS pathway signaling, Brf1 depletion continued to enhance viral replication to a similar extent as observed in control cells (Fig 8C). These results indicate the proviral effect of Brf1 loss is independent of RIG-I/MAVS-mediated interferon signaling and suggests activation of the MAVS pathway does not contribute to the observed antiviral phenotype.

### Brf1 knockdown enhances viral spread of MHV68 in culture

Because our phenotype observed in Brf1-deficient cells was specific to a low-MOI infection and RNA-seq suggested a dysregulated interferon response, we decided to assess for phenotypic changes specific to low MOI infections. High-MOI studies are commonly used to assess phenotypic changes during a single, synchronous replication cycle, whereas low-MOI infections are more complex, as observed phenotypes can arise from differences in viral entry, replication efficiency, or cell-cell spread (Fig. 9A).

**Figure 9.**
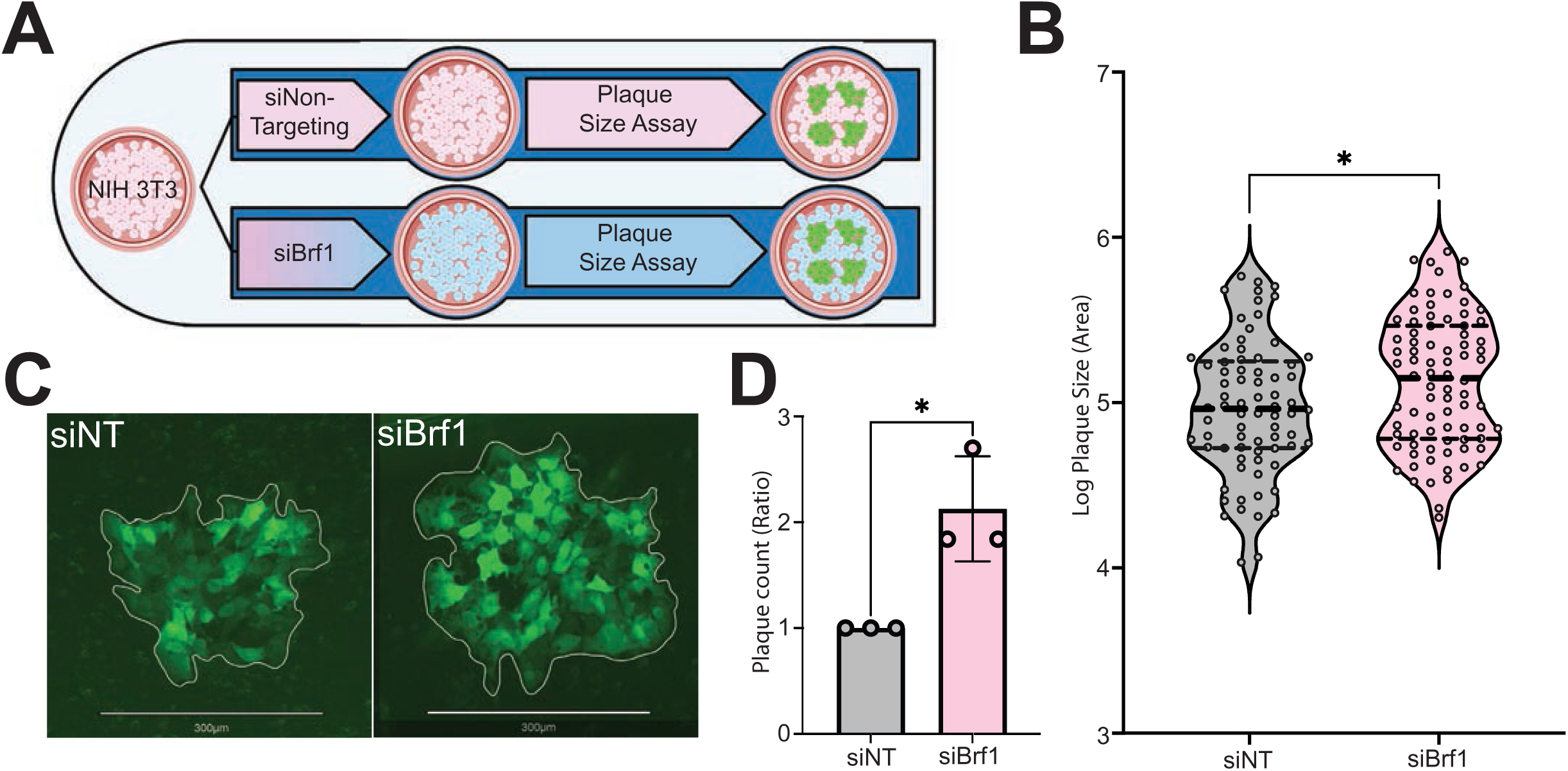
Brf1 knockdown enhances MHV68 plaque size and formation. (A) Schematic of the plaque assay workflow. NIH 3T3 cells were transfected with either non-targeting (siNT) or Brf1-targeting (siBrf1) siRNA, infected with MHV68-GFP at a low MOI, and overlaid with 1% methylcellulose-containing medium to permit plaque formation. (B) Quantification of plaque size, expressed as log-transformed plaque pixel area. (C) Representative images of plaques from siNT-and siBrf1-treated cells, illustrating the average plaque size observed for each condition. (D) Total plaque numbers in siBrf1-treated cells normalized to the corresponding siNT control condition. Statistical significance was determined by one-way ANOVA. P ≤ 0.05 (*), P ≤ 0.01 (**), P ≤ 0.001 (***), P ≤ 0.0001 (****)

Based on prior imaging that revealed altered infection patterns (Fig 4D), as well as a noticeable increase in titer (Fig 4A) produced from Brf1-deficient cultures, we hypothesized that enhanced viral spread may contribute to the observed phenotype. To confirm this, we performed plaque size assays on NIH3T3 control and Brf1-deficient cells, overlaying infected monolayers with methylcellulose to restrict virion diffusion and assess cell-cell spread (Fig. 9A). Brf1-deficient cells exhibited significantly larger average plaque sizes compared with control cells (Fig. 9B), and representative images illustrate this enhanced viral spread in culture (Fig. 9C). Further, we counted the total number of plaques produced on both siNT and siBrf1 monolayer treated with the same viral inoculum. Across three biological replicates, we repeatedly observed 2-fold more plaques on Brf1-depleted monolayers, suggesting that this phenotype, at least in part, is driven by enhanced MHV68 entry (Fig 9D). These findings indicate that Brf1-dependent restriction of entry and/or viral spread, in addition to the dysregulation observed in interferon signaling, contributes to the antiviral phenotype observed under low-MOI conditions.

## Discussion

Although increased activity of RNA polymerase III is a commonly observed phenotype during herpesvirus infection, little is known about why this phenomenon occurs and how it affects viral infection. This study sheds light on this observation by revealing an anti-viral role on Brf1 dependent RNA polymerase III activity during MHV68 infection. It is known that during MHV68 infection, increased RNA polymerase III activity is specifically seen through increases in premature-tRNAs over the course of infection^17^. From this information, we decided to target this synthesis as directly as possible when limiting Pol III activity in our system. Brf1 presented as the best target for knockdown as it is known to be the limiting factor of the transcription factor IIIB complex and is not involved in other cellular processes like the two other protein in the complex, Tbp and Bdp1 ^51–53^. Brf1 knockdown also allows for sustained function of Pol III unlike specific inhibitors that block Pol III function completely which are known to decrease cell viability. While Pol III inhibitors do not have a significant effect on titers in fibroblasts^24^, knocking down Brf1 specifically enhances titer, likely due to specific antiviral activity afforded by activity at Brf1-dependent Pol III promoters.

We observed that the effect of the Brf1 knockdown on Pol III transcripts levels is dependent on the native levels and turnover rate of each transcript. Two premature-tRNAs, Tyrosine and Leucine, showed minimal expression from MHV68 infection in Brf1-deficient cells. Other Brf1 dependent transcripts —5S rRNA and 7SL — showed no difference in gene expression at any point during MHV68 infection (Fig 2A). These results are not unexpected as both 5S and 7SL are natively expressed at about 10,000 times the amount of our pre-tRNAs (Fig 2B). Any changes due to loss of Pol III activity over the course of a three-day infection are overshadowed by high endogenous gene expression enhanced by the inherent stability of these transcripts ^54, 55^. We expect nascent RNA techniques would be required to see the effect of Brf1 knockdown on these transcripts.

We found that the MHV68 viral gene expression, protein accumulation, and virion production are increased during a low MOI MHV68 infection in Brf1-deficient fibroblasts. Interestingly, when performing these same experiments at a high MOI we found no phenotype associated with Brf1 depletion. This suggests that a replication specific phenotype is not at play, and Brf1’s role can influence viral entry or spread, events uniquely measured in a low MOI setting and heavily influenced by an interferon response in the culture. Importantly, our genetic rescue experiments suggest that the role of Brf1 in restricting MHV68 replication is specific and is not due to off-target effects of siRNA knockdown. In these experiments, however, we found that overexpression of Brf1 on its own also caused increases in MHV68 viral titers. The mechanism behind this phenotype is most likely separate from the anti-viral phenotype seen in our Brf1 knockdown experiments. We predict that this phenotype can be attributed to enhanced translation supported by Brf1-dependent transcripts, and this should be dissected in future studies.

One way in which Brf1 could modulate infection is through the production of specific Brf1-dependent Pol III transcripts. However, our attempt at screening the effect of several Brf1-dependent Pol III transcripts on MHV68 infection did not reveal a transcript with anti-viral activity (Fig S1). Surprisingly, we saw a subtle proviral effect following overexpression of our Brf1-independent control transcript, 7SK. While 7SK overexpression led to slight increase in viral titer (did not reach statistical significance), it accomplished this with reduced extracellular genome copies, suggesting an overall increase in the infectivity of virus particles (particle:PFU) (Fig S1E). We observed the reciprocal effect when depleting 7SK in the system. We saw minimal change in viral titer but a modest increase in extracellular genome copies, showing a decrease in infectivity of virus particles (Fig S1G-K). In fact, elevated levels of 7SK were observed at 3 dpi following Brf1 knockdown (Fig 2A), allowing the possibility that 7SK might contribute to the Brf1-dependent phenotype. However, given that 7SK transcript levels only increase towards the endpoint of infection, it is unlikely that 7SK is the main driver of Brf1 antiviral activity and should be explored separately.

To assess the downstream mechanism behind viral enhancement upon Brf1 depletion, RNA sequencing revealed a more rapid interferon induction when Brf1 is knocked down compared to control cells during infection. In Brf1-depleted cells infected at low MOI, interferon responsive genes reached peak expression by 1 dpi, a time when a minority of the cells are infected, as compared to 2 dpi when Brf1 is present. A recent preprint exploring Brf1 knockdown in macrophages indicates that Pol III machinery is situated upstream of immune and inflammatory genes, and loss of Brf1 enhances their expression during MHV68 infection^56^.This observation is highly aligned with the expression data reported here in fibroblasts, as we see a similar enhancement of interferon responsive and inflammation gene expression. We additionally find that this can have important consequences on viral cell to cell spread that is independent of MAVS. Whether Brf1 antiviral activity is dependent on other triggers of interferon-responsive or inflammatory gene pathways will be explored in future studies.

Finally, we are intrigued by the fact that the enhanced interferon-responsive gene expression triggered early by Brf1 depletion can be leveraged by MHV68 through host-shutoff, as we observed a global enhancement of host gene depletion (including IFN responsive genes) in Brf1-deficient fibroblasts at days 2-3 post infection. Viral host-shutoff mechanisms are widely recognized as immune-evasion strategies that suppress Type I interferon production and interferon-stimulated gene expression, and our results are consistent with these studies. The observed drop in interferon-responsive gene expression later in infection following Brf1 depletion likely directly supports infectious virion production and spread. MHV68 cell-cell spread has not been studied in depth, but several interferon-responsive genes, influenced by Brf1 would be predicted to limit MHV68 replication directly or mount antiviral programs in uninfected bystander cells ^57–59^. Future studies should aim to identify the Brf1-dependent factor that drives enhances cell-cell spread, as this may reveal new knowledge regarding cell-cell spread of herpesviruses as well as highlight novel antiviral targets and pathways.

**Figure 10. Graphical abstract: Brf1 dependent RNA Polymerase III antiviral model.** Schematic representation showing Brf1 dependent functional RNA Pol III activity (TOP) vs limited RNA Pol III activity (BOTTOM) and its role in affecting MHV68 viral spread.

**Figure S1. RNA Pol III transcript overexpression and 7SK anti-sense oligonucleotide knockdown.** (A) Relative endogenous expression levels of the analyzed non-coding RNAs measured by RT-qPCR normalized to Pre-tRNA-leu. (B) Schematic showing all the individual transcripts overexpressed by transfection with their corresponding M11 promoter. Red = Type I transcript; Blue = Type II transcript; Green = Type III transcript (C) RT-qPCR showing steady state transcript gene expression levels comparing each individual transcript to endogenous transcript levels in the pUC19 transfected samples. (D) TCID_50_ assays measuring viral titers of MHV68 at 3 dpi, labeled with corresponding overexpression vector. (E) Extracellular genome copies comparing control to 7SK overexpression. (F) MHV68 Particle/PFU measurement of 7SK overexpression. (G) Schematic showing antisense oligonucleotides (ASO) targeting the 5’ end and 3’ end of 7SK transcripts for degradation. (H) RT-qPCR measuring 7SK gene expression at 5 hours post-transfection and 3 days post-infection. Scrambled ASO used as control. (I) TCID_50_ assay showing MHV68 viral titers labeled by corresponding ASO. (J) Extracellular genome copies measured by qPCR labeled by corresponding ASO. (K) Ratio of viral particles to plaque forming units labeled by corresponding ASO. Statistical significance was determined by one-way ANOVA. P ≤ 0.05 (*), P ≤ 0.01 (**), P ≤ 0.001 (***), P ≤ 0.0001 (****)

**Figure S2. RNA Sequencing heat maps and validation.** (A) Heat map of total viral genes separated by kinetic classes. Top portion shows ERCC normalized read counts. Bottom portion shows Log_2_FC comparing Brf1-deficient cells to control cells. (B) Isolated graphs showing mRNA levels of ORF50 and gB from 1-3 dpi. (C) Pathway analysis of 0 dpi showing highly dysregulated pathways based on P_adj_ value and set size. (D) Heat map of interferon related genes that were dysregulated at 1 dpi. Top portion shows ERCC normalized read counts. Bottom portion shows Log_2_FC comparing Brf1-deficient cells to control cells. (E) RT-qPCR measuring gene expression of two upregulated genes from the RNA sequencing dataset, Irgm1 and Irgm2. N=2. Statistical significance was determined by one-way ANOVA. P ≤ 0.05 (*), P ≤ 0.01 (**), P ≤ 0.001 (***), P ≤ 0.0001 (****)

## Acknowledgements

We thank Seqinfomics, LLC for providing bioinformatic support in the development and maintenance of the RNA-seq analysis pipeline. We thank Craig Forrest at the University of Arkansas for Medical Sciences for reagents and manuscript feedback. We thank Rich Roller and his lab at the University of Iowa for project and manuscript feedback. This research was funded by the National Institutes of Health, grant R21AI173703 to JMT; grant T32AI007533 to KR; and the Office of Undergraduate Research at the University of Iowa, ICRU fellowship to ZG; a Diversity in Cancer Research (DICR) Post Baccalaureate Grant # DICR POST-BACC-22-1041993-01-DPBACC to CH; NSF REU DBI-2244169 to RE and CH.

